# The FraA adhesin and Ebp pili mediate *Enterococcus faecalis* adhesion under fluid flow

**DOI:** 10.64898/2026.11.12.766216

**Authors:** Haris Antypas, Mélanie Roch, Kenneth Kok Fei Ng, Brenda Silva Rosa da Luz, Rachel Jing Wen Tan, Riya Zita Thomas, Alain Filloux, Kimberly A. Kline

**Affiliations:** Singapore Centre for Environmental Life Sciences Engineering, School of Biological Sciences, Nanyang Technological University, Singapore, Singapore; MRC Centre for Molecular Bacteriology and Infection, Department of Life Sciences, Imperial College London, London, United Kingdom; Department of Microbiology and Molecular Medicine, University of Geneva, Geneva, Switzerland

## Abstract

*Enterococcus faecalis* is a frequent cause of infective endocarditis and catheter-associated urinary tract infections (CAUTI). Successful colonization is challenged by blood and urine flow and relies on a strong and resilient bacterial attachment. Here, we identify molecular determinants that enable *E. faecalis* to maintain adhesion under flow, notably the flow-related adhesin (FraA), a previously uncharacterized adhesion factor encoded by *OG1RF_11764*. Transcriptomic analysis revealed increased *fraA* expression in surface-attached bacteria exposed to shear stress. Structural predictions identified an N-terminal region with homology to characterized Gram-positive adhesins, followed by tandem CLIPPER repeats with putative isopeptide-bond-forming residues and a C-terminal cell-wall sorting signal. Using microfluidics and single-cell tracking, we found that deletion of *fraA* accelerated detachment from hydrophobic abiotic surfaces under fluid flow. In line with this finding, loss of *fraA* reduced catheter colonization in a murine CAUTI model. Analysis of 9,192 *E. faecalis* genomes identified FraA in 50.8% of isolates, spanning clinical and non-clinical sources, with sequence type-associated variation in CLIPPER repeat number. Another determinant, Ebp pilus, enhances adhesion at high shear stress and contributes to a shear-enhanced adhesion pattern. Collectively, these findings identify FraA as an adhesion factor contributing to medical-device colonization and together with Ebp pili support attachment across different hydrodynamic conditions. Identifying these adhesion mechanisms could inform strategies to prevent colonization of medical devices exposed to fluid flow.

## INTRODUCTION

Bacterial pathogens encounter host niches that differ in physicochemical cues, and their ability to establish infection requires adaptation to these local microenvironments ^1,2^. One such cue is fluid flow, a defining feature of the vascular system and urinary tract. Blood circulation and urine flow provide a first line of defence by imposing hydrodynamic forces that challenge bacterial attachment, thereby limiting colonization. Despite these forces, these host niches remain susceptible to pathogens that express virulence determinants enabling attachment under flow, leading to infections such as infective endocarditis (IE) and urinary tract infections (UTI), which are associated with substantial morbidity and healthcare costs ^3,4^. Moreover, medical interventions introduce additional surfaces for bacterial colonization, including urinary catheters, vascular catheters, and prosthetic valves. Infections associated with these devices impose a considerable burden on healthcare systems ^5^. Understanding how bacteria establish and maintain attachment under flow is therefore central to understanding pathogenesis in these host niches and identifying strategies to prevent infection by targeting adhesion.

Bacterial adhesion under fluid flow is mediated by surface appendages, such as pili, and non-pilus-associated adhesins. On biotic surfaces, these bind host receptors or macromolecules and support attachment against shear stress ^6^. Examples include *Escherichia coli* P fimbriae, whose tip adhesin PapG recognizes Galα1–4Gal moieties in globoseries glycolipids on renal epithelial cells ^7–9^, supporting attachment under flow in nephrons ^10,11^. *Neisseria meningitidis* uses type IV pili to adhere to vascular endothelial cells under flow ^12^, while *Staphylococcus aureus* exploits von Willebrand factor-binding protein to attach to the vascular wall under shear stress ^13^. In some cases, increasing shear stress enhances bacterial adhesion. In *E. coli*, the type 1 fimbriae tip adhesin FimH binds mannose-containing receptors on host cells, including uroplakin Ia on bladder epithelial cells^14,15^. FimH forms catch bonds, which exhibit shear stress-enhanced adhesion, thereby strengthening attachment under flow ^16,17^. Shear-enhanced adhesion also occurs in Gram-positive bacteria. *Streptococcus gordonii* serine-rich repeat adhesins bind sialylated glycans on the platelet receptor GPIbα and promote platelet attachment as shear increases ^18^.

Beyond specific ligand-receptor interactions, bacteria can attach to and colonize abiotic surfaces, such as implants and catheters, through binding to adsorbed host proteins ^19^ or direct interactions with the underlying material, governed by hydrophobic, electrostatic, and van der Waals interactions ^20^. For example, *Acinetobacter baumannii* Csu pili promote adhesion to polystyrene, polypropylene, and polyethylene through hydrophobic finger-like loops in the tip adhesin Csu^21^. The *Pseudomonas fluorescens* adhesin LapA supports attachment to both hydrophilic and hydrophobic surfaces ^22^, while the *Staphylococcus epidermidis* surface protein SdrF promotes adhesion to Dacron and polystyrene ^23^. Despite the exposure of vascular and urinary catheters to blood and urine flow, respectively, the mechanisms supporting bacterial attachment to abiotic surfaces under shear stress remain less well characterized. In *Pseudomonas aeruginosa*, type IV pilus retraction tilts cells away from abiotic surfaces at low shear stress, promoting detachment. Higher shear stress counteracts this tilting, thereby enhancing adhesion ^24^. This demonstrates that shear-enhanced adhesion can also arise through the mechanical interplay between surface appendages and hydrodynamic forces on abiotic surfaces.

*Enterococcus faecalis* is a Gram-positive opportunistic pathogen, frequently isolated from infections defined by fluid flow, such as infective endocarditis (IE), central line-associated bloodstream infections (CLABSI), and catheter-associated urinary tract infections (CAUTI). It is the fifth most common pathogen isolated from adult healthcare-associated infections in the US, accounting for 7.6% and 7.7% of pathogens reported from CLABSIs in hospital wards and intensive care units, respectively, and 12.1% of CLABSI pathogens in long-term acute-care hospitals^25^. *E. faecalis* also accounts for approximately 6.5-11.7% and 11.1% of CAUTI infection in the US and Europe, respectively ^25,26^. Its frequent recovery from these infections is consistent with its ability to adhere to both host extracellular matrix components and abiotic medical device surfaces. The adhesin to collagen of *E. faecalis* (Ace) binds type I and IV collagen and laminin and contributes to IE-associated adhesion ^27,28^, whereas the endocarditis and biofilm-associated (Ebp) pili bind fibrinogen and platelets and contribute to adhesion in IE and CAUTI, with the metal ion-dependent adhesion site of the tip adhesin EbpA mediating colonization during CAUTI ^19,29–32^. Additional proteins involved include the *E. faecalis* surface proteins Fss1–3, which bind fibrinogen, and enterococcal fibronectin-binding protein A (EfbA), which binds fibronectin and contributes to adhesion and pathogenesis in IE and urinary tract infection^33–35^. *E. faecalis* also expresses factors implicated in adhesion to abiotic surfaces. The pCF10-encoded PrgA mediates adhesion and biofilm formation on abiotic surfaces ^36^ and the enterococcal surface protein Esp promotes biofilm formation on polystyrene and bacterial deposition on bile-drain materials under flow ^37,38^. Consistent with this adhesion repertoire, *E. faecalis* has been reported to colonize both glass and fibrinogen-coated surfaces more efficiently under high-flow than low-flow conditions, attributed to its ability to form linear chains of cells^39^. Nevertheless, the molecular determinants that enable attachment to withstand shear stress are not defined.

We recently demonstrated that exposing surface-attached *E. faecalis* to fluid flow mimicking the shear stress experienced by aortic valve vegetations in IE upregulates 59 genes and downregulates 167 genes relative to static conditions (|log_2_FC| > 1.0, FDR < 0.05) ^40^. We found that fluid flow suppresses Fsr quorum sensing, which in turn promotes biofilm formation and worsens IE outcomes. Here, we used this transcriptomic dataset to investigate how fluid flow influences adhesion factor expression and identified a previously uncharacterized adhesin with increased expression under flow, which we named flow-related <u>a</u>dhesin A (FraA). We show that FraA enhances adhesion under flow and contributes to catheter colonization *in vivo*, while Ebp pili further support the ability of *E. faecalis* to withstand shear stress. Collectively, these findings identify FraA and Ebp pili as determinants of adhesion under flow and link FraA to the colonization of medical devices.

## RESULTS

### Identification of a novel adhesion factor flow-related adhesin (FraA) expressed by surface-attached *E. faecalis* under shear stress

To discover molecular determinants that enable *E. faecalis* to withstand shear stress on abiotic surfaces, we revisited our previous transcriptomic dataset^40^ comparing gene expression in OG1RF WT attached under physiologically relevant shear stress (20 dynes/cm^2^) or static conditions to a hydrophobic flow chamber surface (**Fig. S1**). We shortlisted 21 genes encoding proteins that could be present at the cell surface. These contain the tripartite pattern required for cell wall sorting, i.e. a C-terminal LPXTG-family motif, a hydrophobic transmembrane tail, and a positively charged cytoplasmic terminus^41^. Among these 21 proteins, 16 had a predicted N-terminal Sec signal peptide with SignalP 6.0 ^42^ and are thus candidate sortase-anchored surface proteins (**Table S1**). Of these 16 genes, only 2 showed increased expression under shear stress (**Fig. 1A**): *ace* (log_2_FC +1.73, FDR = 0.0018) and a previously uncharacterized gene *OG1RF_11764/OG1RF_RS13215* (log_2_FC +2.10, FDR = 0.0072), which we named flow-related <u>a</u>dhesin <u>A</u> (*fraA*). OG1RF_12251, encoding a small LPXTG protein, was downregulated (log_2_FC −1.54, FDR = 0.019). The remaining 13 genes were not differentially expressed, including the previously identified pilus genes *ebpABC* (log_2_FC +0.02 to +0.07, FDR ≥ 0.79), the fibrinogen-binding adhesin gene *fss1* (log_2_FC +0.19, FDR = 0.50), and *elrB* (log_2_FC −1.06, FDR = 0.47), a member of the recently characterized *elrABCDE* pilus assembly operon (**Fig. 1A; Table S1**) ^43^.

**Fig. 1.**
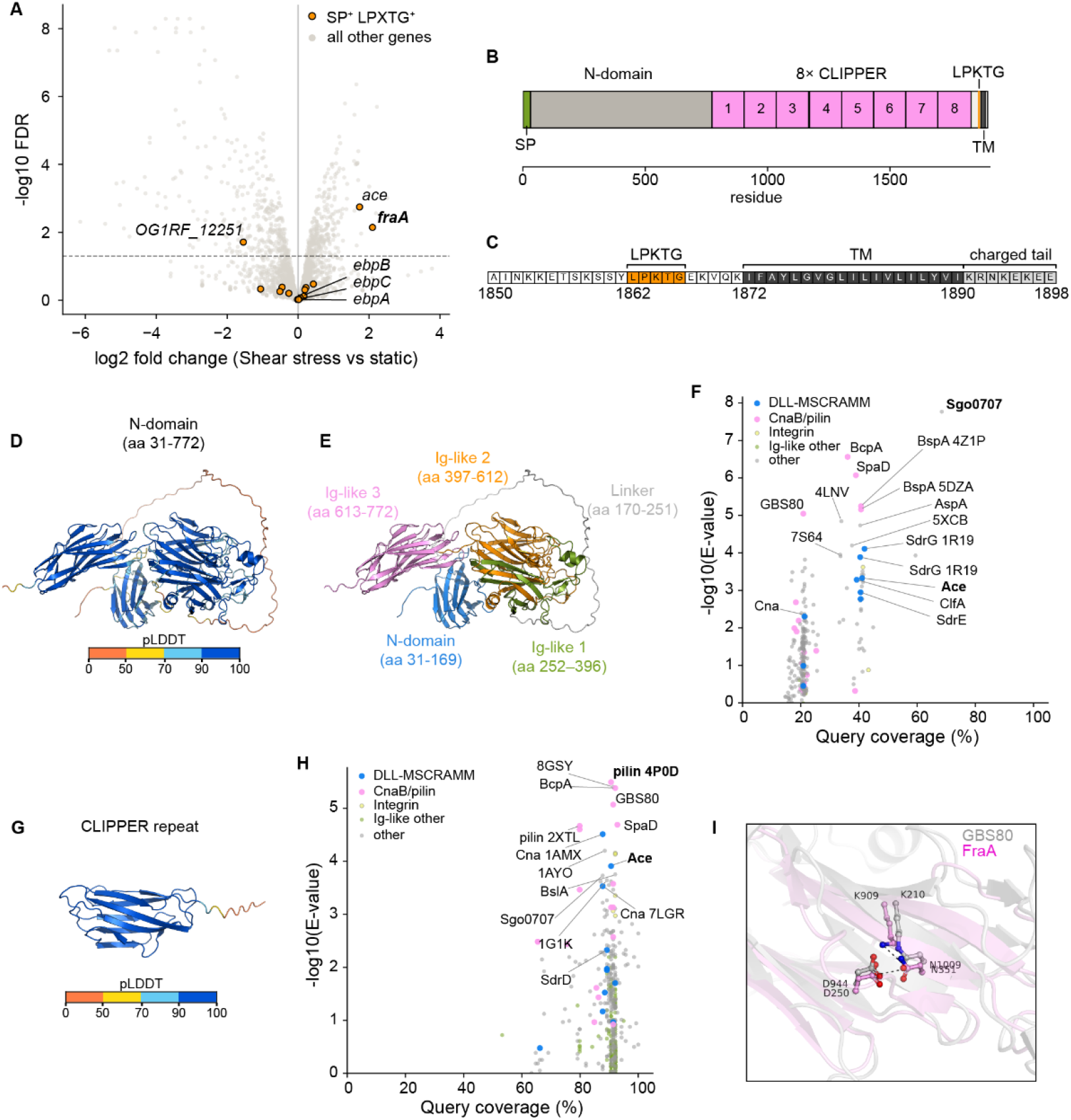
Identification of a novel adhesion factor flow-related adhesin (FraA) expressed by surface-attached *E. faecalis* under shear stress. **A.** Volcano plot of differential gene expression in shear stress-exposed attached *E. faecalis* OG1RF compared to attached bacteria under static conditions. N = 3. Gray = all 2,583 detected genes, orange = 16 predicted signal peptide positive (SP^+^) LPXTG sortase substrates. Dashed line indicates FDR = 0.05. **B.** Domain architecture of FraA (OG1RF_11764). The 1,898-aa protein is drawn to scale. Signal peptide (SP, green), N-terminal domain (gray, aa 31–772), eight tandem CLIPPER repeats (pink, numbered 1–8; ∼131 aa each), LPKTG sorting motif (orange, aa 1862–1866), and transmembrane helix (TM; dark gray, aa 1872– 1890). **C.** C-terminal inset (aa 1840–1898) showing the LPKTG motif, transmembrane helix, and charged tail (KRNKEKEE) that constitute the cell-wall sorting signal. **D.** ColabFold predicted structural model for the N-terminal domain (aa 31–772) of FraA (mean pLDDT 84.5). **E.** ColabFold structural model for the N-terminal domain with color coded predicted domains. The N-terminal region (aa 31–169; mean pLDDT 79.4) contains 10 β-strands arranged into two opposing β-sheets and is followed by a low-confidence linker and three putative Ig-like domains with mean pLDDT values of 92.4, 93.2 and 95.6, respectively. **F.** Foldseek search of the ColabFold model of the FraA N-terminal domain against PDB100. Each point represents one PDB chain hit, plotted by query coverage and −log10(E-value) and colored by structural family: DLL-MSCRAMM (blue), CnaB/pilin (pink), integrin (green), other Ig-like (olive) and other folds (gray). **G.** ColabFold predicted structural model for a single CLIPPER repeat (aa 904– 1042) of FraA (mean pLDDT 93.9). **H.** Foldseek search of the ColabFold model of the single CLIPPER repeat unit against PDB100. Each point represents one PDB chain hit, plotted by query coverage and −log10(E-value) and colored by structural family: DLL-MSCRAMM (blue), CnaB/pilin (pink), integrin (green), other Ig-like (olive) and other folds (gray). **I.** Superposition of the FraA CLIPPER repeat model (pink) on the GBS80 pilin domain (PDB 3PF2, chain A, gray; CEalign RMSD 3.54 Å over 128 residues). FraA residues Lys909, Asp944 and Asn1009 align with the isopeptide-linked Lys210 and Asn351 and catalytic Asp250 of GBS80, with corresponding atom distances of 1.77, 0.88, 1.18 Å. Dashed lines indicate the Lys909-NZ···Asn1009-CG and Asp944-CG···Asn1009-CG distances in the FraA model.

Next, we investigated whether the predicted domains of FraA supported a role in adhesion. Sequence analysis revealed an N-terminal Sec signal peptide (**Fig. 1B**, **Table S1**), followed by a ∼740-residue N-terminal domain, 8 repeats of a Cross-Linked IsoPeptide Protein in the Extracellular Region (CLIPPER) domain ^44^ (**Fig. 1B**, **Fig. S2**; formerly DUF11 domains), and a C-terminal cell-wall sorting signal (**Fig. 1B-C**). We modelled the N-terminal domain and a representative CLIPPER repeat to examine their structural similarity to characterized proteins. ColabFold models of the N-terminal domain (aa 31–772) and a single CLIPPER repeat (aa 904–1042) were predicted with high confidence (**Fig. 1D, G**). The N-terminal model contained a β-sandwich region followed by three putative Ig-like domains (**Fig. 1E, Figs. S2–S3**). A Foldseek search against PDB100 identified the N-terminal domain of the collagen-binding *Streptococcus gordonii* adhesin Sgo0707 as the highest-ranking structural match (**Fig. 1F, Table S2**) ^45^. The collagen-binding A domain of *E. faecalis* Ace also partly matched the FraA N-terminal domain (**Fig. 1F, Table S2**). Foldseek analysis of a single CLIPPER repeat identified similarities to Ig-like domains from Gram-positive pilins and MSCRAMMs, including the T6 backbone pilin of *Streptococcus pyogenes* and *E. faecalis* Ace (**Fig. 1H, Table S3**). Consistent with the presence of intramolecular isopeptide bonds in CLIPPER domains, Lys909, Asp944 and Asn1009 in the FraA model adopted a geometry compatible with isopeptide bond formation (**Fig. 1I**; pink). These residues aligned closely with the experimentally resolved isopeptide-forming residues and catalytic aspartate of the *Streptococcus agalactiae* GBS80 pilin (**Fig. 1I**; grey), suggesting that the FraA repeats could also form an isopeptide-stabilized region. Collectively, these results support FraA as a candidate adhesin.

### Expression of *fraA* enhances adhesion under shear stress

To investigate whether *fraA* plays a role in adhesion under shear stress, we constructed a deletion mutant Δ*fraA* (**Table S4**). WT and Δ*fraA* cells were allowed to attach in a microfluidic chamber and then exposed to a stepwise increase of shear stress from 1 to 80 dyn/cm^2^, which falls within a physiological range encountered in the human body ^46–48^. We tracked individual bacteria by time-lapse microscopy and measured how long they remained attached. Detachment between Δ*fraA* and WT was compared using a Cox proportional-hazards model. The hazard ratio (HR) describes how likely Δ*fraA* bacteria are to detach at any given time relative to WT, with HR > 1 indicating faster detachment, HR < 1 indicating slower detachment, and HR = 1 indicating no difference. Immediately after introducing flow exerting 1 dyn/cm^2^ shear stress, Δ*fraA* adhesion rapidly decreased compared to WT (**Fig. 2A-B**; HR > 1). Detachment of Δ*fraA* was faster compared to WT from 1 to 40 dyn/cm², although this difference was gradually attenuated with increasing shear stress. By the end of the 40-dyn/cm² step only ∼21% of Δ*fraA* cells remained attached versus ∼43% of WT. After being exposed to 80 dyn/cm², both strains converged to ∼11–13% adhesion at the endpoint, suggesting that this shear stress may approach the limits of attachment for most cells under these experimental conditions, irrespective of FraA. When we quantified adhesion of Δ*fraA* under static conditions, we observed no significant difference compared to WT (**Fig. 2C**). These data indicate that *fraA* enhances attachment to abiotic surfaces under shear stress, consistent with its increased expression under flow and its predicted adhesin function.

**Fig. 2.**
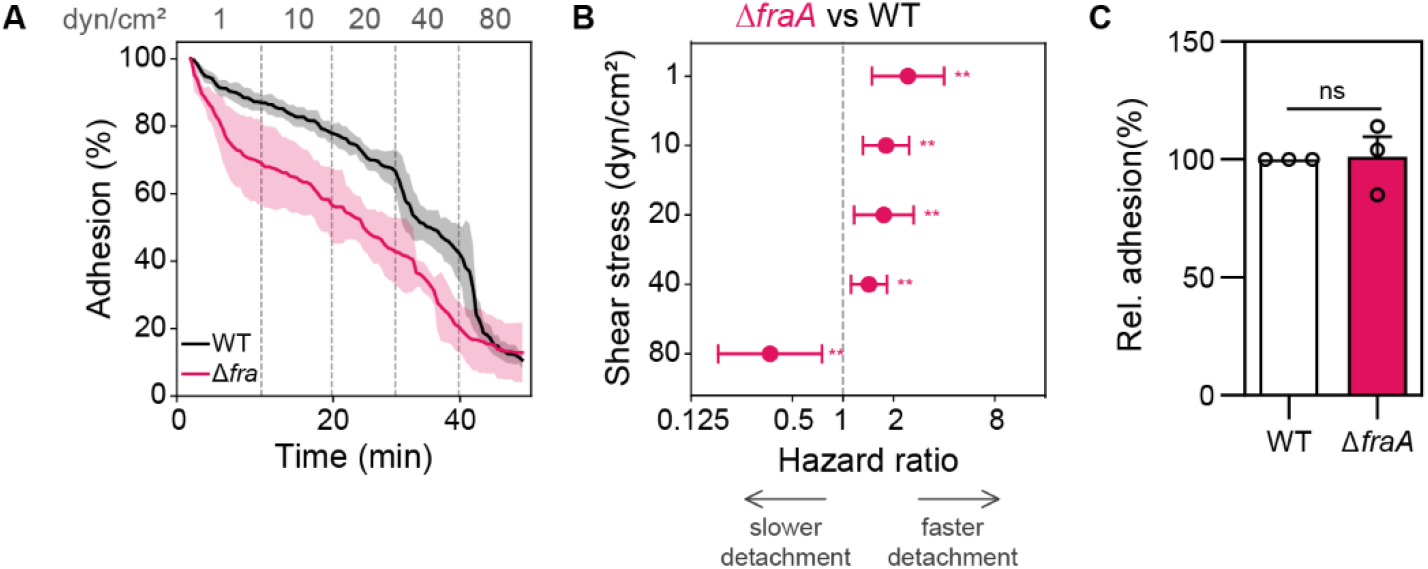
Expression of *fraA* enhances adhesion under shear stress. **A.** Bacterial adhesion (%) over a stepwise shear stress increase. Dashed lines mark transitions between 1, 10, 20, 40 and 80 dyn/cm². Adhesion is normalized to the total number of bacteria on the first time-lapse frame at t=0. Mean ± SEM is shown, N = 3. **B.** Cox proportional-hazards analysis of detachment hazard ratios (HR) for Δ*fraA* vs WT at each shear step with 95% CIs. Risk set at each shear stress step corresponds to bacteria still attached at step start. Asterisks denote Benjamini–Hochberg adjusted P values (**P < 0.01). HR > 1 indicates faster detachment than WT; HR < 1 indicates slower detachment. **C.** Bacterial adhesion (%) of Δ*fraA* relative to WT under static conditions on a microfluidic channel. Statistical significance was assessed with one sample t-test. Mean ± SEM is shown, t = 15 min, N = 3, ns = not significant.

### Expression of *fraA* contributes to catheter colonization *in vivo*

*E. faecalis* is one of the most common pathogens isolated from CAUTI, in which bacteria have to colonize the abiotic surface of the catheter under flow of urine from bladder emptying. To investigate whether *fraA* mediates adhesion in CAUTI, we implanted a silicone catheter piece in the bladder of mice and infected with WT or Δ*fraA E. faecalis*. We found that catheter colonization was reduced in the absence of *fraA* compared to WT (**Fig. 3A**, p < 0.05), whereas no significant difference was observed in bladder and kidney CFU (**Fig. 3B, C**). These data further support a role for FraA in mediating adhesion to medical devices implanted in host niches exposed to hydrodynamic forces.

**Fig. 3.**
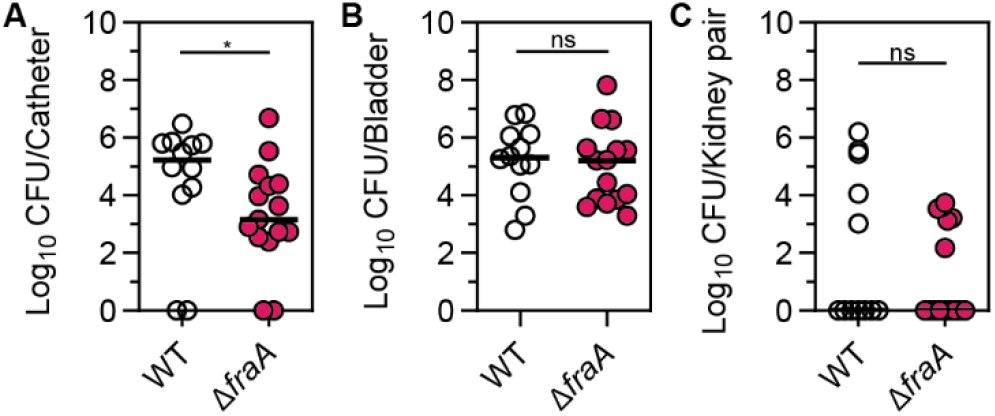
Expression of *fraA* contributes to catheter colonization *in vivo*. **A-C.** Median CFU in catheter (A), bladder (B) and kidney pairs (C) harvested from a murine CAUTI model at 24 h post-infection. 3-6 mice per group per experiment from 3 independent experiments. Statistical significance was assessed with a Mann-Whitney test. * = p < 0.05, ns = not significant

### FraA is variable in length and is encoded by both clinical and non-clinical *E. faecalis* isolates

To investigate the distribution of *fraA* across *E. faecalis*, we analysed 9,192 *E. faecalis* genome sequences from the NCBI Pathogen Detection database, representing isolates classified as clinical, non-clinical, or unknown sources. We first identified candidate *fraA* loci by searching the genome assemblies against the OG1RF FraA protein sequence using TBLASTN, then predicted protein-coding sequences within the matching genomic regions. Because the CLIPPER repeat number in FraA varies across genomes, our minimum requirement for identifying full-length FraA was that its predicted protein product contained both the first N-terminal subdomain (aa 40-168) and the cell wall sorting signal (aa 1862-1898). Using this criterion, 4,672 genomes (50.8 %) encoded full-length FraA. Partial FraA were classified separately according to whether the incomplete sequence was contained within a single contig or interrupted at a contig edge, consistent with assembly fragmentation. Full-length FraA was detected in both 2,024 clinical isolates (52.1%) and 1,293 non-clinical isolates (45.5%) (**Fig. 4A**). Among clinical isolates, its prevalence was broadly similar across isolation sources, including gastrointestinal/fecal (55.1%), bloodstream (57.4%), urinary tract (49.7%), and skin, wound, or soft tissue isolates (55.2%) (**Fig. 4B**).

**Fig. 4.**
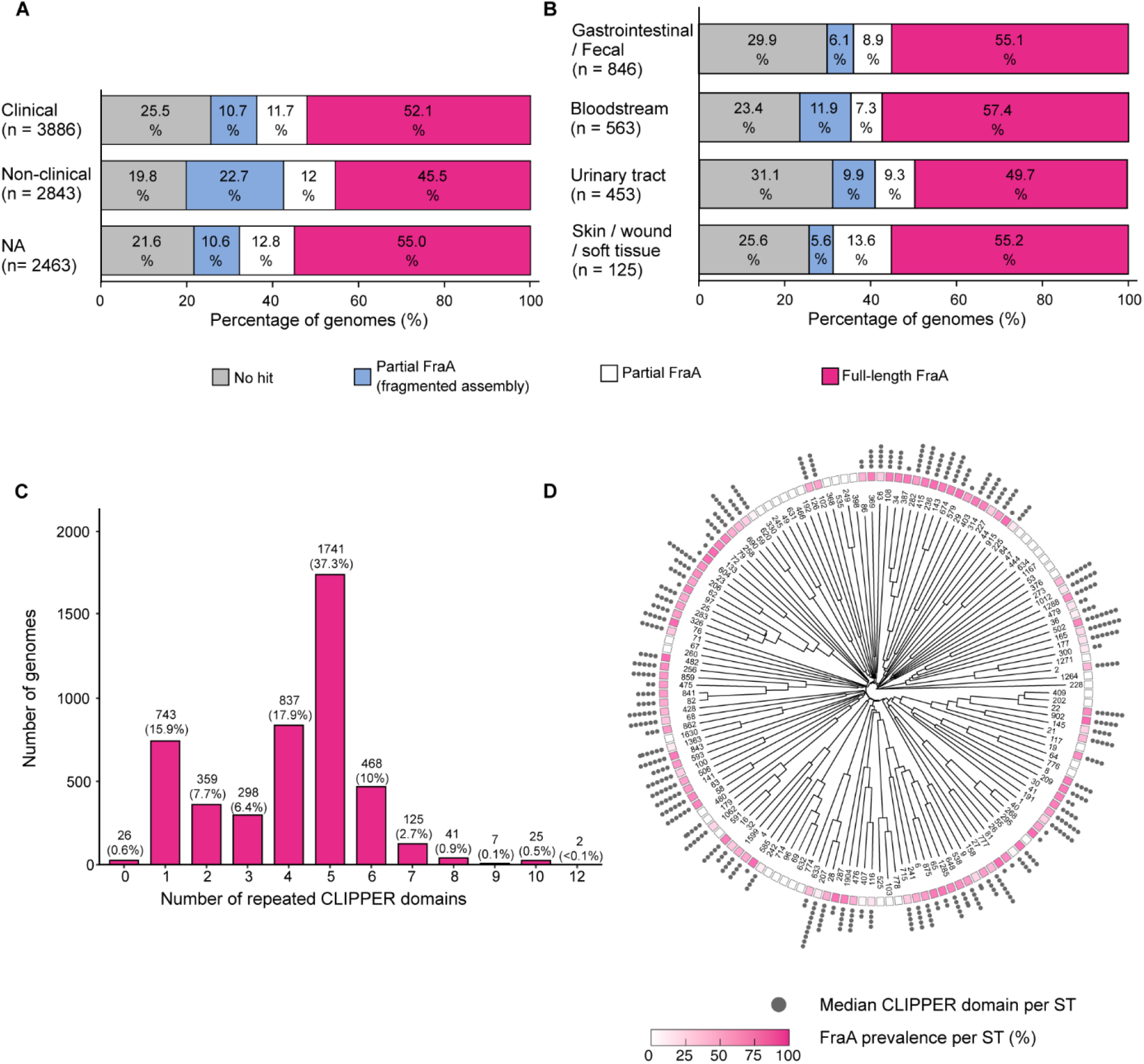
FraA is variable in length and is encoded by both clinical and non-clinical *E. faecalis* isolates. (A) Distribution of FraA protein categories among isolates classified as clinical, non-clinical, or having unknown isolation source (NA). Full-length proteins (magenta) were defined by the presence of the N-terminal region (aa 40-168) through the cell-wall sorting signal (aa 1862-1898). Partial FraA were classified separately according to whether the incomplete sequence was contained within a single contig or interrupted at a contig edge, consistent with assembly fragmentation. Genomes without a detectable hit are shown in gray. Numbers of genomes are indicated in parentheses, and percentages are shown within bars. (B) Distribution of the same categories among clinical isolates, stratified by isolation source: gastrointestinal/fecal, bloodstream, urinary tract, and skin/wound/soft tissue. (C) Distribution of CLIPPER repeat-domain copy numbers among full-length FraA proteins (n = 4,672). Labels indicate the number and percentage of genomes in each category. (D) Distribution of full-length FraA prevalence and median CLIPPER repeat-domain copy number across sequence types (STs). The neighbor-joining tree was constructed from cgMLST allele distances using one representative genome per ST with ST numbers indicated at the tips. The colored ring indicates the percentage of genomes encoding full-length FraA within each ST. Outer gray dot chains represent the median number of CLIPPER repeat domains among full-length FraA proteins within each ST, with each dot representing one domain.

Among isolates carrying full-length FraA, the predicted proteins contained between 0 and 12 CLIPPER domains, with 5 copies being the most frequent configuration (37.3%; **Fig. 4C**). Proteins containing 7 or more copies accounted for only 4.3% of full-length hits. PCR analysis of selected isolates supported the observed repeat-number variation as a genuine biological variation rather than a genome assembly artifact. (**Fig. S4**). The number of CLIPPER domains varied significantly across sequence types (ST; Kruskal– Wallis χ² = 2518.2, df = 74, p < 2.2 × 10⁻¹⁶) with a large effect size (ε² = 0.717), indicating strong lineage-associated conservation of protein length (**Fig. S5**). To determine whether variation in FraA prevalence and CLIPPER repeat number was restricted to particular genetic lineages, we mapped full-length FraA prevalence and median CLIPPER domain copy number onto a neighbor-joining tree derived from cgMLST allele distances for 148 STs (**Fig. 4D**). STs with low or undetectable full-length FraA prevalence occurred in several distinct branches of the cgMLST-based tree, suggesting that FraA may have been independently lost across different lineages. Together, these findings show that full-length FraA is broadly distributed among clinical and non-clinical *E. faecalis* isolates and that CLIPPER repeat number is strongly associated with ST.

### Ebp pili enhance adhesion of *E. faecalis* to abiotic surfaces under high shear stress

Adhesion in the absence of *fraA* was attenuated but not completely abrogated under flow, indicating that other surface-associated factors may be involved. To that end, we tested whether Ebp pili contribute to adhesion under flow. We exposed WT and mutants lacking the entire pilus (Δ*ebpABC*) or only the tip adhesin (Δ*ebpA*) to uniform fluid flow conditions exerting 1, 10, 20, 40, or 80 dynes/cm^2^. WT exhibited catch-bond behaviour, with slower detachment at 20 dyn/cm² than 1 dyn/cm² (**Fig. 5A, B**; HR < 1). Deletion of either *ebpA* or *ebpABC* abolished this behaviour (**Fig. 5A, C**). At high shear stress (80 dyn/cm²) both Ebp pilus mutants detached faster than WT (**Fig. 5A, C**; HR > 1). Together, these findings suggest that Ebp-piliated bacteria are better equipped to withstand high shear stress, whereas bacteria lacking Ebp pili remain well adapted to low shear conditions.

**Fig. 5.**
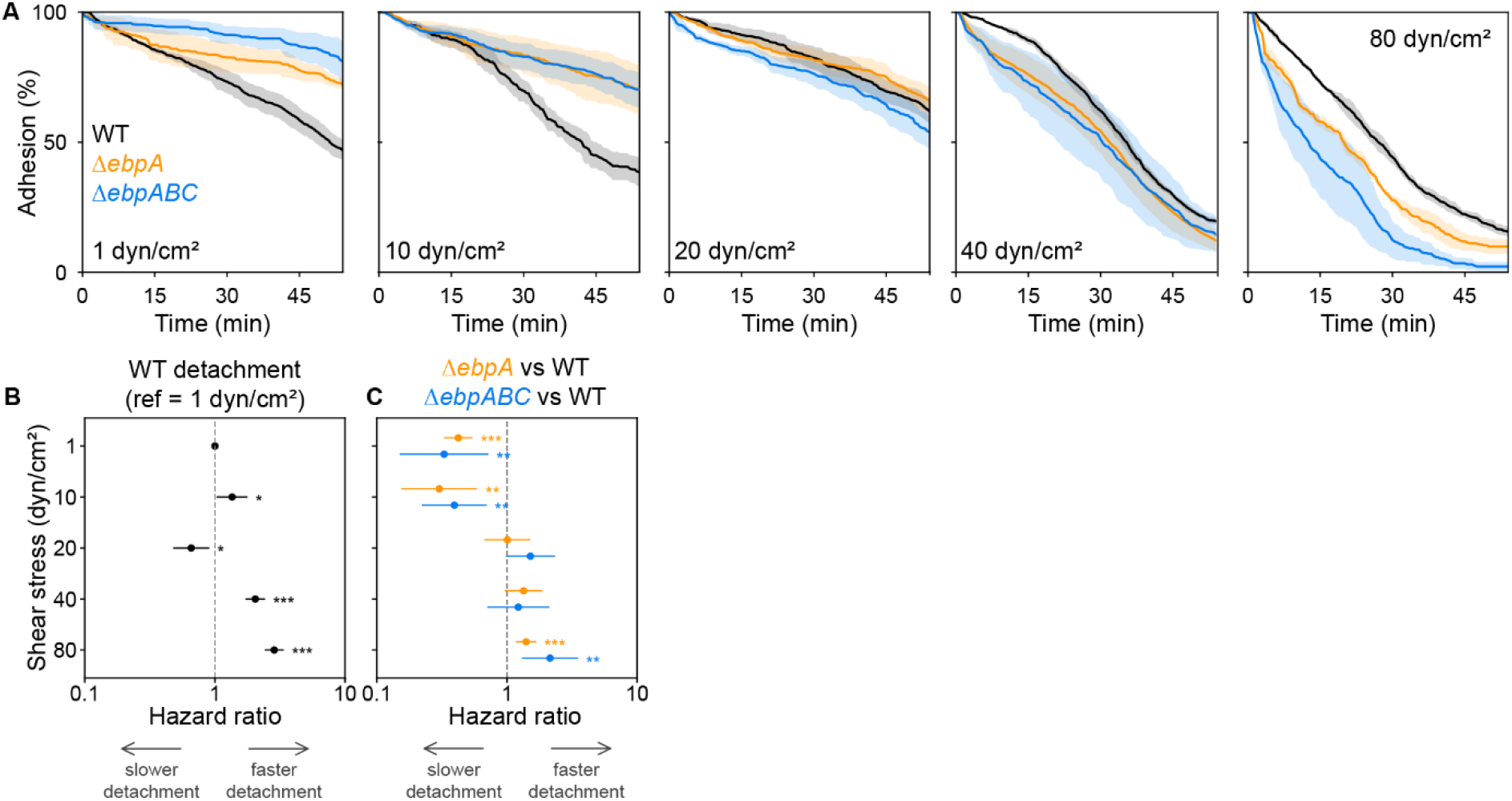
Ebp pili enhance adhesion of *E. faecalis* to abiotic surfaces under high shear stress. **A.** Adhesion of *E. faecalis* OG1RF wild-type (WT, black), Δ*ebpA* (orange) and Δ*ebpABC* (blue) under fluid flow at the indicated shear stresses (1, 10, 20, 40 and 80 dyn/cm²). Bacteria were allowed to attach for 10 min under static conditions and then exposed to flow for 60 min. Adhesion at each time point is expressed as a percentage of the initial (t = 0) value for each replicate. Curves show the mean and the shaded band the SEM across replicate videos. **B-C**. Hazard ratios (HR) from Cox proportional-hazards models comparing WT detachment at each shear stress with that at 1 dyn/cm² (B) and detachment of Δ*ebpA* (orange) and Δ*ebpABC* (blue) with WT at the same shear stress (C). Points show HRs and horizontal lines indicate 95% confidence intervals. HR > 1 indicates faster detachment and HR < 1 indicates slower detachment relative to the respective reference. WT at 1 dyn/cm² in B is shown at HR = 1 without a confidence interval. Asterisks denote Benjamini–Hochberg-adjusted P values (*P < 0.05, **P < 0.01, ***P < 0.001); N = 3 independent experiments per strain and shear stress combination.

## DISCUSSION

The mechanical forces exerted by urine flow and blood circulation provide a first line of defence against bacterial colonization of the urinary tract and vasculature. Yet *E. faecalis* colonizes these niches, causing infections such as CAUTI and IE. The adhesin repertoire that enables *E. faecalis* to withstand fluid flow has not been identified, as enterococcal adhesins have been largely characterized for their ability to bind host ligands under static conditions. Here, we used a microfluidic model to challenge *E. faecalis* attachment to abiotic surfaces across a range of physiologically relevant shear stress values. We identified FraA, a previously uncharacterized adhesin, and Ebp pili as determinants that enhance attachment under shear stress. FraA contributed at low shear stress, whereas Ebp pili supported attachment at high shear stress and contributed to a shear-enhanced adhesion pattern. Loss of FraA also reduced catheter colonization in a murine CAUTI model, establishing its relevance to medical-device colonization. Together, these findings reveal how different adhesins support *E. faecalis* attachment across the range of shear stress encountered on medical devices exposed to blood or urine flow, providing insight into the mechanisms underlying their colonization.

FraA exhibits structural similarities to several proteins with experimentally established roles in adhesion. Notably, the Sgo0707 N-terminal domain, the strongest match to the FraA N-terminal region, contributes to *Streptococcus gordonii* adhesion to collagen-coated surfaces under flow ^45^. Part of the FraA N-terminal region also shares structural similarity to the N1–N2 ligand-binding region of the *E. faecalis* collagen adhesin Ace^49^. Whether the FraA N-terminal domain mediates adhesion remains to be experimentally tested. The variation in CLIPPER repeat number suggests that FraA may belong to the broader class of periscope proteins ^50^, whose variable-length repeat regions control the projection of functional domains from the bacterial surface. It is proposed that repeat-number variation enables bacteria to modify surface interactions in response to selection pressure, potentially influencing host colonization, biofilm formation and immune evasion ^50^. Longer stalks may increase the accessibility of adhesive domains, whereas shorter variants may reduce their exposure to immune recognition. Changes in protein length may also influence access to neighbouring surface proteins. Similarly, variation in FraA repeat number could alter the presentation of its N-terminal domain and its interactions with surrounding surfaces. Whether these differences affect attachment under flow or recognition by the host immune system remains to be determined. Importantly, complete *fraA* deletion reduced but did not abolish adhesion under flow. Together with the presence of *fraA* in 50.8% of isolate genomes analysed and its lack of a strong association with the urinary tract, this finding suggests that FraA may be one of several determinants that mediate adhesion under flow. We hypothesize that FraA-negative strains maintain adhesion through other surface factors, including the widely conserved Ebp pili and Ace adhesin ^30,51^, with their relative contributions depending on the surface and flow conditions encountered.

Our findings show that *E. faecalis* exhibits shear-enhanced adhesion to hydrophobic surfaces, with slower detachment at intermediate shear stress than at low or high shear stress. Loss of Ebp pili abolished this response, resulting in a slip-bond-like adhesion pattern. Whether this behaviour reflects force-dependent changes in adhesin binding, cell orientation, or access of other surface factors to the substrate remains to be determined. A plausible interpretation is that pili change the geometry of cell–surface contact. In *P. aeruginosa*, type IV pilus retraction promotes cell tilting and departure at low shear, whereas increasing shear favours horizontal attachment and prolongs adhesion^24^. Although Ebp pili lack this retraction mechanism, asymmetric distribution on newly divided cells^52^ could influence cell orientation during attachment, limiting close cell–surface contact or access by other adhesins at low shear. The latter is also supported by previous findings that Ebp pili inhibit aggregation-substance-mediated cell–cell interactions ^53^. Loss of the EbpA tip adhesin also abolished the shear-enhanced pattern of adhesion. Although this suggests that EbpA might mediate this catch-bond adhesion pattern, we cannot exclude that the enhanced adhesion of Δ*ebpA* at low shear stress may also reflect changes in pilus expression, as deletion of *ebpA* has been reported to produce fewer pili ^54^. Further investigation using atomic force microscopy– based single-molecule force spectroscopy could determine whether Ebp pili or EbpA alone mediates shear-enhanced adhesion.

Although *fraA* expression was higher in surface-attached bacteria exposed to flow, our experimental design cannot establish whether shear stress induces its expression. Because transcriptomic analysis was restricted to bacteria that remained attached, the flow-exposed population was enriched for cells able to withstand shear stress. Higher *fraA* expression could therefore reflect preferential retention of bacteria with pre-existing elevated expression, transcriptional induction in response to flow, or a combination of both. Future studies using *fraA* promoter–reporter constructs to track expression in individual bacteria before and during flow exposure could distinguish gene upregulation from enrichment for cells with higher *fraA* expression.

Consistent with its ability to colonize host sites exposed to fluid flow, *E. faecalis* exhibited a remarkable ability to maintain adhesion under prolonged exposure to fluid flow, even on the planar surfaces of the microfluidic channels without a pre-applied host-protein coating. In the host, surface topography and conditioning films may further support adhesion. For instance, the irregular surface topology of fibrin-rich vegetations could provide shelter from the high flow rates of the heart, while fibrinogen deposits on urinary catheters provide adhesion ligands. While our current *in vivo* findings suggest *fraA* expression in the mouse bladder, further defining where the FraA adhesin is expressed in the host and how it is regulated will increase our understanding of its role in virulence. Nevertheless, its contribution to catheter colonization already highlights its relevance to CAUTI, a major healthcare-associated infection that prolongs hospitalization and increases healthcare costs^55^. The protection afforded by EbpA vaccination and antibody-based therapies in experimental models ^19,56^ further supports targeting enterococcal adhesins to prevent infection. Defining the role of these adhesins under host-relevant shear stress conditions could inform the development and evaluation of such interventions, particularly for medical devices exposed to blood or urine flow.

## METHODS

### Bacterial cultures and construction of deletion mutants

Strains used in this study are shown in **Table S4**. Clinical isolates were collected from patients with wound infection at Tan Tock Seng Hospital, Singapore, under the supervision of T. M. S. Barkham (Laboratory Medicine) and identified as *E. faecalis* strains by MS. All strains were routinely grown in brain heart infusion (BHI) broth at 37 °C without shaking, in ambient air. Primers used to construct deletion mutants in this study are listed in **Table S5**. The OG1RF Δ*11764* mutant was generated by allelic replacement using the temperature-sensitive shuttle vector pGCP213, as previously described ^32,57^. Briefly, pGCP213 was linearized with *EcoRI* and *HindIII* (New England Biolabs, USA), and the corresponding amplified inserts were ligated with T4 DNA ligase (New England Biolabs, USA) and transformed into *E. coli* Stellar competent cells. *E. coli* transformants were selected on LB agar containing 500 µg/mL erythromycin. Plasmid constructs were confirmed by Sanger sequencing and introduced into wild-type OG1RF. Transformants were selected on BHI agar containing 25 µg/mL erythromycin and subsequently passaged at 42°C with erythromycin to promote plasmid integration into the chromosome. Plasmid excision was induced by serial passage at 30°C without antibiotics, and erythromycin-sensitive colonies were screened by PCR for the identification of clean Δ*fraA* mutants.

### Shear stress calculation

We applied shear stresses of 1–80 dyn/cm² to encompass reported estimates for model aortic valve leaflets (3–79 dyn/cm²)^46^ and urethral flow during urination (3–5 dyn/cm²)^47^. Although urinary catheter models have used lower shear stresses (∼0.1 dyn/cm²)^48^, our pump could not reliably generate values below 1 dyn/cm², which therefore defined the lower limit of our experiments. To achieve 1-80 dynes/cm^2^ in microfluidic channels (µ-Slide VI 0.1, ibidi), we calculated the required flow rate using the following formula provided by the manufacturer:

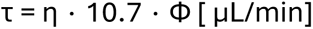

where τ = shear stress (dynes/cm^2^), η = dynamic viscosity (dynes × s/cm^2^) and Φ = flow rate. Dynamic viscosity of BHIS media is 0.0101171 dyn × s/cm^2^ at 37°C, as determined using a viscometer. Thus, shear stresses of 1, 10, 20, 40, and 80 dyn/cm² correspond to flow rates of approximately 9.24, 92.38, 184.75, 369.50, and 739.01 µL/min, respectively.

### Bacterial adhesion assay under shear stress

To quantify bacterial adhesion under flow conditions, uncoated flow chambers (µ-Slide VI 0.1, ibidi) were loaded with overnight cultures of various bacterial strains grown in BHI broth. Chambers were incubated for 10-20 min at 37°C under static conditions to allow for bacterial attachment. Once attachment was established, the chambers were transferred to a microscope stage housed in a 37°C incubator. Continuous flow of BHI supplemented with 40% horse serum (BHIS) was initiated using a syringe pump (KDS Legato 270P, KD Scientific), generating defined shear stress levels of 1, 10, 20, 40 or 80 dyn/cm² for a duration of 50-60 minutes. In a separate experimental setup, shear stress was incrementally increased from 1 to 80 dyn/cm² in 10-minute increments for 50 min. Images were captured every 30 seconds from 1-2 fields of view per microchannel at 40x magnification under phase contrast microscopy. Bacteria were tracked throughout the time-lapse series using ImageJ’s TrackMate plugin. Any bacterial aggregates were excluded. To plot adhesion (%), we calculated the number of attached bacteria per frame and normalized the data to the number of bacteria at t = 0, corresponding to 100 % adhesion.

### Video contact angle measurement

A #1.5 polymer coverslip with an uncoated surface (Cat. No. 10813, ibidi) and the well bottoms of a TC-treated and non-treated polystyrene (Cat. No. 144530, ThermoFisher Scientific) 24-well plate were used. Contact angle measurement was performed using a drop shape analyzer (KRÜSS). 2-4 water drops per surface were tested.

### In silico protein domain analysis

The FraA protein sequence (locus tag OG1RF_11764; GenBank AEA94451.1; RefSeq WP_014525034.1; 1,898 residues) was retrieved from the *E. faecalis* OG1RF genome assembly (NCBI RefSeq assembly GCF_000172575.2)^58^. The N-terminal signal peptide was predicted with SignalP 6.0 ^42^ using the Gram-positive organism type and the Sec/SPI cleavage-site model (predicted cleavage between residues 30 and 31; Sec/SPI likelihood 0.9908). Conserved domains were identified by searching the full-length FraA protein sequence in InterProScan ^59^. The N-terminal head domain (aa 31–772) was defined as the region between the predicted signal-peptide cleavage site and the first CLIPPER repeat. Pairwise sequence identity between the CLIPPER repeat units was computed by global pairwise alignment (BioPython PairwiseAligner; BLOSUM62 substitution matrix; gap-open penalty −10, gap-extension penalty −0.5) as the number of identical positions divided by the total aligned length. A self-similarity dotplot of the full protein (exact 10-mer matches) was generated to visualize the tandem repeat architecture. Structural models of the FraA N-terminal domain (aa 31–772) and a single repeat unit (aa 904–1042) were predicted with ColabFold (v1.6.3)^60^. Each model was searched against the PDB100 database using the Foldseek web server (search.foldseek.com)^61^. Hits were deduplicated by target protein (best E-value retained) and assigned to structural families by PDB title keywords. The isopeptide-bond triad (Lys909, Asp944, Asn1009) was identified in the ColabFold repeat model by geometry (Lys NZ–Asn CG and Asp CG–Asn CG distances) and predicted by CEalign superposition in PyMOL (Schrödinger) onto the crystal structures of GBS80 (PDB 3PF2), which carry experimentally confirmed isopeptide bonds. The corresponding figures were rendered in PyMOL.

### Bacterial adhesion quantification under static conditions

Flow chamber channels (uncoated, ibidi µ-slide VI^0.4^) were seeded with bacterial overnight cultures (∼120 µL) and incubated under static conditions at 37°C for 15 min to allow adhesion. Supernatant was aspirated and channels were washed twice with PBS. Attached bacteria in flow chambers were fixed with 4% paraformaldehyde for 15 min at 37°C and washed twice with PBS. Bacteria were permeabilized with 0.1% Triton in PBS for 5 min, followed by 2 washes with PBS for 5 min each. Bacteria were stained with DAPI (diluted 1:500; stock concentration 5 mg/mL) in PBS for 5 min, followed by a final PBS wash. 10 fields of view distributed along each channel to avoid sampling bias were captured at 40x with fluorescence microscopy. Attached bacteria were enumerated in ImageJ (1.54p) and counts from 10 fields of view (technical replicates) were averaged for each independent experiment (biological replicate). Relative adhesion was calculated by dividing the mean count for Δ*fraA* by the corresponding WT mean within each experiment and multiplying by 100.

### Murine catheter-associated urinary tract infection model

Female C57BL/6J mice 6-7 weeks old were used in accordance with NTU Institutional Animal Care and Use Committee (Protocol A23094). The experimental procedure was carried out as previously described^62^. Briefly, a three-segment catheter was prepared on 30G needles, comprising a 7 mm PE tubing at the shaft, a 5 mm silicone tubing acting as the adaptor, and a 4 mm silicone catheter tubing, to be inserted into the bladder. To prepare the inoculum, *E. faecalis* OG1RF WT and Δ*fraA* strains were incubated overnight in BHI at 37°C, washed in PBS and normalized to OD600 of 0.5 (2-6 × 10^8^ CFU/mL). Anesthesia was induced and maintained with 2% isoflurane. Bladders of anesthetized mice were catheterized via the urethra with the three-segment catheter needle, and the silicone segment was implanted in the bladder by moving forward the PE tubing at the shaft of the needle. A 1-mL syringe with bacterial inoculum was connected to a single segment catheter and used to inoculate 50 µL of bacteria into the mouse slowly. The catheter was slowly retracted, and the animal was returned in its cage. 24 hours post-infection, mice were euthanized with CO_2_ asphyxiation, and the catheter, bladder and kidney pairs were harvested from each mouse to determine bacterial burden. The harvested catheter was placed in sterile 1x PBS on ice, and organs were placed in pre-weighed lysing matrix M tubes on ice. Organs were homogenized 3 times for 20 s at 4 m/s, and catheters sonicated in a water bath at 30% power and 30 kHz for 15 minutes at room temperature. The homogenates were then serially diluted to 10^-5^ in sterile PBS in a 96-well plate and plated on BHI and BHI-rifampicin plates using the drop count method. The plates were incubated overnight at 37°C and CFUs were counted.

### *Enterococcus faecalis* genome dataset and sequence analysis

All *Enterococcus faecalis* genome assemblies available in the NCBI Pathogen Detection database as of June 4, 2026, were considered for analysis. Assemblies were available for 11,763 of the 14,471 isolates listed in the database. Assembly quality was assessed using CheckM2 (v1.1.0), and assemblies with estimated completeness ≥99% and contamination ≤5% were retained. 8 assemblies not assigned to *E. faecalis* were also excluded, yielding a final dataset of 9,192 genomes (**Table S6**). Isolate origin was classified using the associated metadata as clinical, non-clinical (environmental/other), or unknown. Clinical isolates were further categorized by isolation source when this information was available. The main categories were gastrointestinal/fecal, bloodstream, urinary tract, and skin/soft tissue/wound samples. The *fraA* locus (OG1RF_RS13215/OG1RF_11764 in *E. faecalis* OG1RF) was initially identified by TBLASTN searches (BLAST+ v2.17.0+) against the genome assemblies using the OG1RF FraA protein sequence as the query. Matching genomic regions were extracted with up to 5 kb of flanking sequence on either side using SAMtools (v1.13). Protein-coding sequences within these regions were predicted using Prodigal v2.6.3 to ensure consistent annotation across genomes. Predicted proteins were searched using BLASTP, with OG1RF FraA sequences corresponding to the 3 N-terminal query regions (aa 40-168, 265-607, and 613-764), the cell wall sorting signal domain (aa 1862-1898), and a representative CLIPPER repeat domain (aa 773-903) as queries. Domain hits were retained when both amino acid sequence identity and query coverage exceeded 90%.

A predicted FraA protein was classified as full-length when both the first N-terminal subdomain and the cell wall sorting signal domain were identified within a single Prodigal-predicted protein. Incomplete FraA were classified as partial proteins contained within a single contig or as partial proteins associated with fragmented assemblies when the sequence was interrupted at a contig edge. Genomes without a qualifying hit were classified as no hit. CLIPPER domain copy number was determined by counting retained repeat-domain BLASTP hits within each full-length FraA protein. Full-length FraA prevalence was calculated as the percentage of genomes meeting the full-length detection criterion within each isolate-origin category, clinical isolation-source category, or sequence type (ST).

To visualize the distribution of FraA across genetic lineages, STs represented by at least 10 genomes in the final dataset were retained irrespective of FraA status, yielding 148 STs. One representative genome per ST was included in the core-genome multilocus sequence typing (cgMLST) analysis. An allele-distance matrix was generated using Ridom SeqSphere+ (v12.07.7) with the predefined *E. faecalis* 1,972-target cgMLST scheme. Missing values were ignored in pairwise distance calculations. A neighbor-joining tree was inferred from this matrix using the nj function in the R package ape, midpoint rooted and displayed in a circular layout. External annotations showed the prevalence of full-length FraA and the median CLIPPER domain copy number among full-length FraA proteins within each ST.

### PCR detection of *fraA* length variability

Colony PCR was used to assess variation in *fraA* length among *E. faecalis* strains using primers 11764_FW and 11764_RV (**Table S5**) and Ǫ5 High-Fidelity DNA Polymerase. Individual colonies were resuspended in 50 µL of Milli-Ǫ water, boiled for 5 min and cooled on ice. 1 µL of lysate was used per 25-µL reaction. PCR products were resolved on 0.8% agarose gels in TBE buffer alongside a GeneRuler 1 kb DNA ladder.

### Statistical analysis

To determine whether detachment differences under shear stress were statistically significant, we applied Cox proportional-hazards regression, a survival-analysis method that estimates the relative rate of bacterial detachment over time. The result is a hazard ratio (HR): an HR > 1 indicates faster detachment, whereas an HR < 1 indicates slower detachment relative to the reference condition. Because no re-attachment was observed during exposure to fluid flow, the per-frame count of attached bacteria remained stable or decreased, and the count at each time point directly gave the number of bacteria that had not yet detached. Thus, each decrease in bacteria between two consecutive frames represents one or more detachment events during that 30 s interval, and all events in that interval were assigned a detachment time equal to the later frame. Bacteria still attached at the endpoint were treated as right-censored. Each analyzed time-lapse recording represented an independent bacterial culture and experiment, with 3 biological replicates per strain and shear stress condition. Cluster-robust standard errors were calculated with time-lapse recording as the clustering unit. When shear stress was incrementally increased, we fitted a separate Cox model at each shear stress step, each comparing Δ*fraA* with WT and using only the bacteria still attached at the start of that step. For the constant-shear experiments, WT detachment was compared across shear stresses using 1 dyn/cm² as the reference (Fig. 5B). At each shear stress, Δ*ebpA* and Δ*ebpABC* were compared with WT in a model containing all three strains (Fig. 5C). P values were corrected with the Benjamini–Hochberg procedure. Adjusted P < 0.05 was considered significant. All Cox regression analyses used lifelines v0.30 in Python. Statistical significance comparing mean adhesion (%) under static adhesion was assessed with one sample t-test and median CFU per catheter or tissue with a Mann-Whitney test. Differences in CLIPPER domain copy number among STs were assessed using a Kruskal-Wallis test in R, restricted to the 75 STs represented by at least 10 genomes encoding full-length FraA. The magnitude of the association between ST and repeat number was quantified using the rank-based effect size epsilon-squared (ε²) in R, with larger values indicating greater differences among STs.

## FUNDING

Funding for this work was provided by the Singapore National Research Foundation and Ministry of Education Singapore under its Research Centre of Excellence Program (SCELSE). This study was supported by the Wallenberg Foundation Postdoctoral Fellowship at Nanyang Technological University Singapore, SCELSE Seed Funding (SF-05), Open Fund – Young Individual Research Grant by the National Medical Research Council Singapore (MOH-000939), all to H.A. This work was also supported by the Singapore Ministry of Education under its tier 2 program (MOE2019-T2-2-089) and by a Swiss National Science Foundation (SNSF) project grant (219227), as well as a Professor Installation grant from the Société Académique de Genève, all awarded to K.A.K.

## ACKNOWLEDGEMENTS

We thank Jazelyn Lew Shi Ǫi (SCELSE) and Ronni Anderson Gonçalves da Silva (SMART) for assistance with preliminary experiments, and Caroline Stéphanie Manzano (UNIGE) for assistance with cloning.

## AUTHOR CONTRIBUTIONS

Conceptualization: H.A., K.A.K.

Methodology: H.A., M.R., K.A.K.

Formal Analysis: H.A., M.R., K.A.K.

Investigation: H.A., M.R., K.K.F.N., B.S.R.D.L., R.J.W.T., R.Z.T.

Resources: H.A., A.F., K.A.K.

Writing - Original draft preparation: H.A.

Writing-Review and editing: H.A., M.R., K.K.F.N., B.S.R.D.L., R.J.W.T., R.Z.T., A.F., K.A.K.

Visualization: H.A., M.R.

Supervision: H.A., K.A.K.

Project administration: H.A., K.A.K.

Funding Acquisition: H.A., K.A.K.

## COMPETING INTERESTS

The authors declare no competing interests.

## DATA AVAILABILITY

Transcriptomic data reanalyzed in this study have been deposited in NCBI under the bioproject accession numbers PRJNA1219810 and PRJNA1219807. All publicly available genomes used to investigate FraA frequency are listed in Table S6.

## Notes

### Competing Interest Statement

The authors have declared no competing interest.

