## Supplementary Materials for "The FraA adhesin and Ebp pili mediate *Enterococcus faecalis* adhesion under fluid flow"

### Contents

**Fig. S1.** Hydrophobicity of surfaces used in this study

**Fig. S2.** Internal sequence similarity and conservation of FraA CLIPPER repeats

**Fig. S3.** Predicted secondary-structure topology of the FraA CLIPPER repeat and N-terminal domain

**Fig. S4.** Variation in CLIPPER domain number among *E. faecalis* strains

**Fig. S5.** FraA CLIPPER domain number across *E. faecalis* sequence types

**Table S4.** Bacterial strains used in this study

**Table S5.** Primers used in this study

References

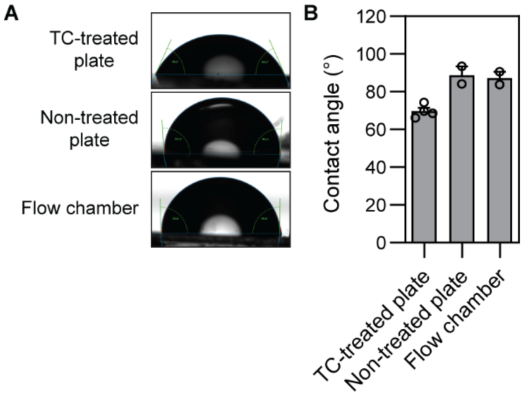


**Supplementary Fig. S1. Hydrophobicity of surfaces used in this study. A-B.** Flow chamber surface hydrophobicity is comparable to a non-treated well plate. Representative water-droplet images (A) and contact angles for a tissue-culture-treated (TC-treated) plate, a non-treated plate, and the flow-chamber surface (B) assessed by video contact-angle measurement are shown. Higher contact angles indicate more hydrophobic surfaces. The average of 2-4 repeat measurements from N = 1 is shown. Error = SEM.

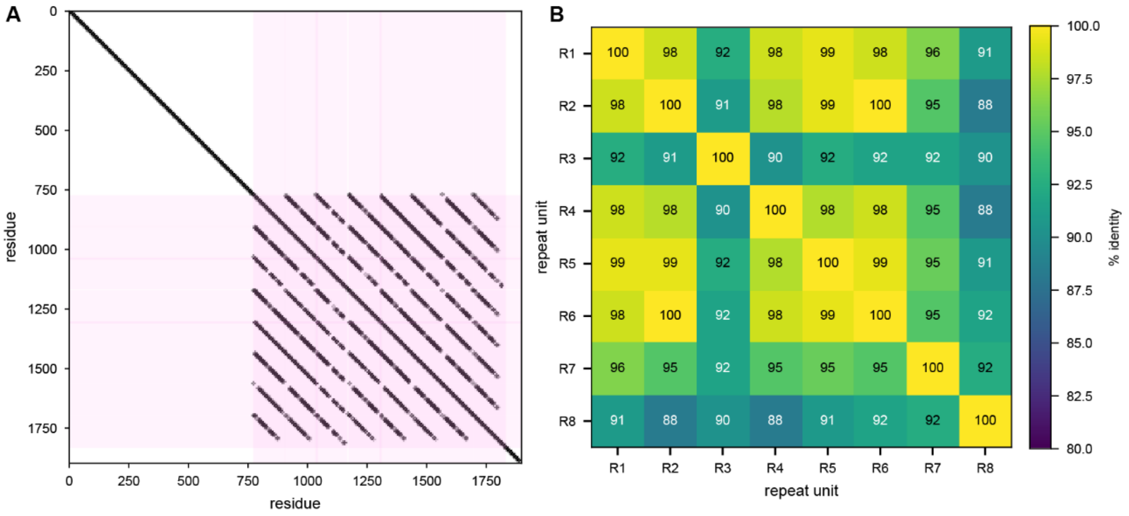


**Supplementary Fig. S2. Internal sequence similarity and conservation of the tandem CLIPPER repeats in FraA. A.** Self-similarity dot plot of the full-length FraA (1,898 aa), generated using exact 10-aa matches. The main diagonal represents identity of the sequence with itself. Pink shading marks the region containing eight tandem CLIPPER repeat units, spanning approximately residues 773–1832. Parallel off-diagonal lines within this region result from sequence matches between different repeat units. No comparable internal repeats were detected elsewhere in the protein using this criterion. **B.** Pairwise amino acid sequence identity among repeat units R1–R8, calculated from global pairwise alignments scored using BLOSUM62. Values denote percentage identity. Excluding self-comparisons along the diagonal, pairwise identity ranged from 88% to 100%.

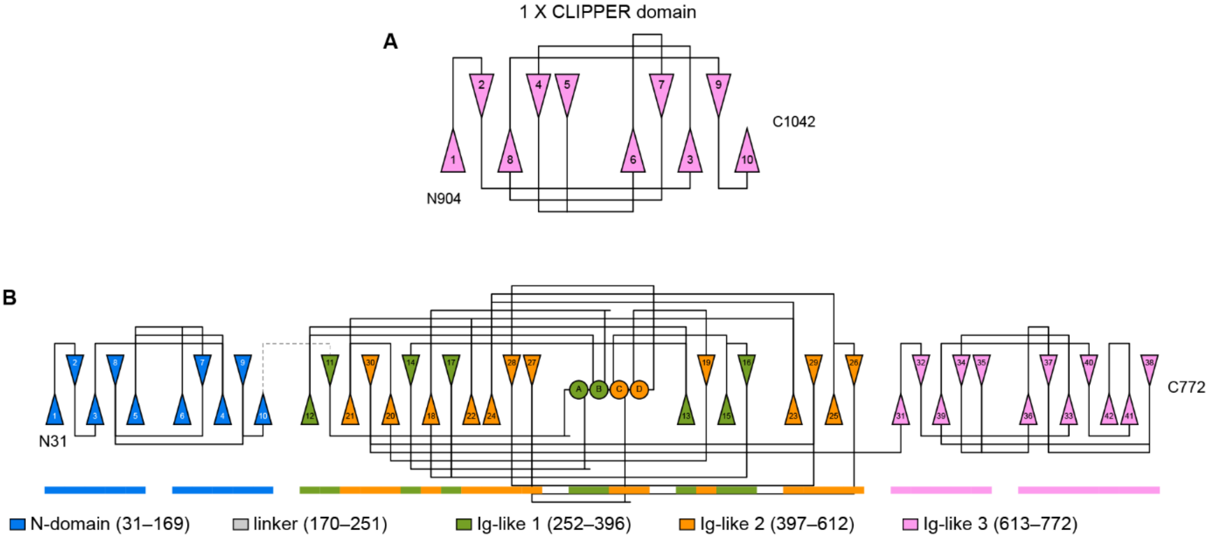


**Supplementary Fig. S3. Predicted secondary-structure topology of the FraA CLIPPER repeat domain and N-domain. A-B.** The topology diagram was generated from the highest-ranked ColabFold model of FraA residues 904-1042 for the CLIPPER domain (A) and 31–772 for the N-domain (B). Analysis of the CLIPPER domain predicted 10 β-strands (triangles) and no helices (circles) arranged in 2 opposing β-sheets . Analysis of the N-domain model predicted 4 structured regions separated by an 82-aa low-confidence segment. The N-terminal domain (residues 31–169aa; blue) comprised ten β-strands arranged into two opposing β-sheets. Residues 170–251 contained no assigned regular secondary structure and had a mean pLDDT of 29.4 (linker, dashed grey line). The remainder of the model comprised three putative Ig-like domains: Ig-like domain 1 (green, 252–396aa), Ig-like domain 2 (orange, 397–612aa), and Ig-like domain 3 (pink, 613–772aa). These domains contained 7, 13, and 12 β-strands, respectively. Four short α-helices were confined to the first 2 Ig-like domains, whereas the N-terminal domain and third Ig-like domain were entirely β-stranded. The 3 Ig-like domains were predicted with high local confidence, with mean pLDDT values of 92.4, 93.2 and 95.6, compared with 79.4 for the N-terminal domain.

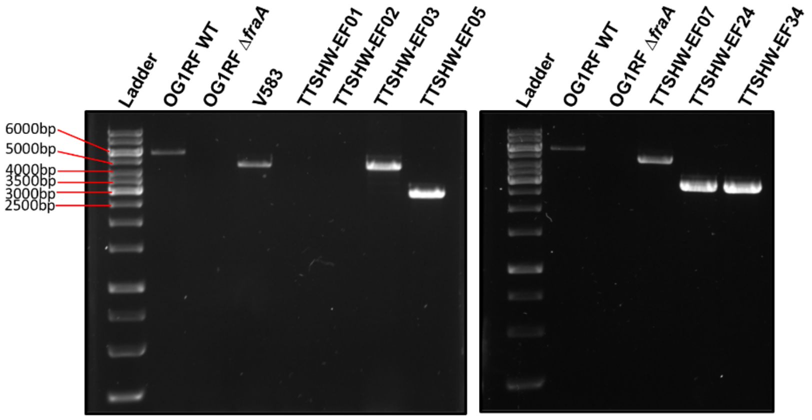


| Strain | CLIPPER domains (n) |
|---|---|
| OG1RF | 8 |
| V583 | 5 |
| TTSHW-EF01 | *fraA*⁻ |
| TTSHW-EF02 | *fraA*⁻ |
| TTSHW-EF03 | 5 |
| TTSHW-EF05 | 2 |
| TTSHW-EF07 | 5 |
| TTSHW-EF24 | 2 |
| TTSHW-EF34 | 2 |

**Supplementary Figure S4. CLIPPER domain number in *fraA* varies across *E. faecalis* strains.** PCR amplification of the *fraA* ORF in *E. faecalis* strains with variable CLIPPER repeat number or complete absence of *fraA (fraA⁻)*. Gel electrophoresis results and corresponding strain table are shown.

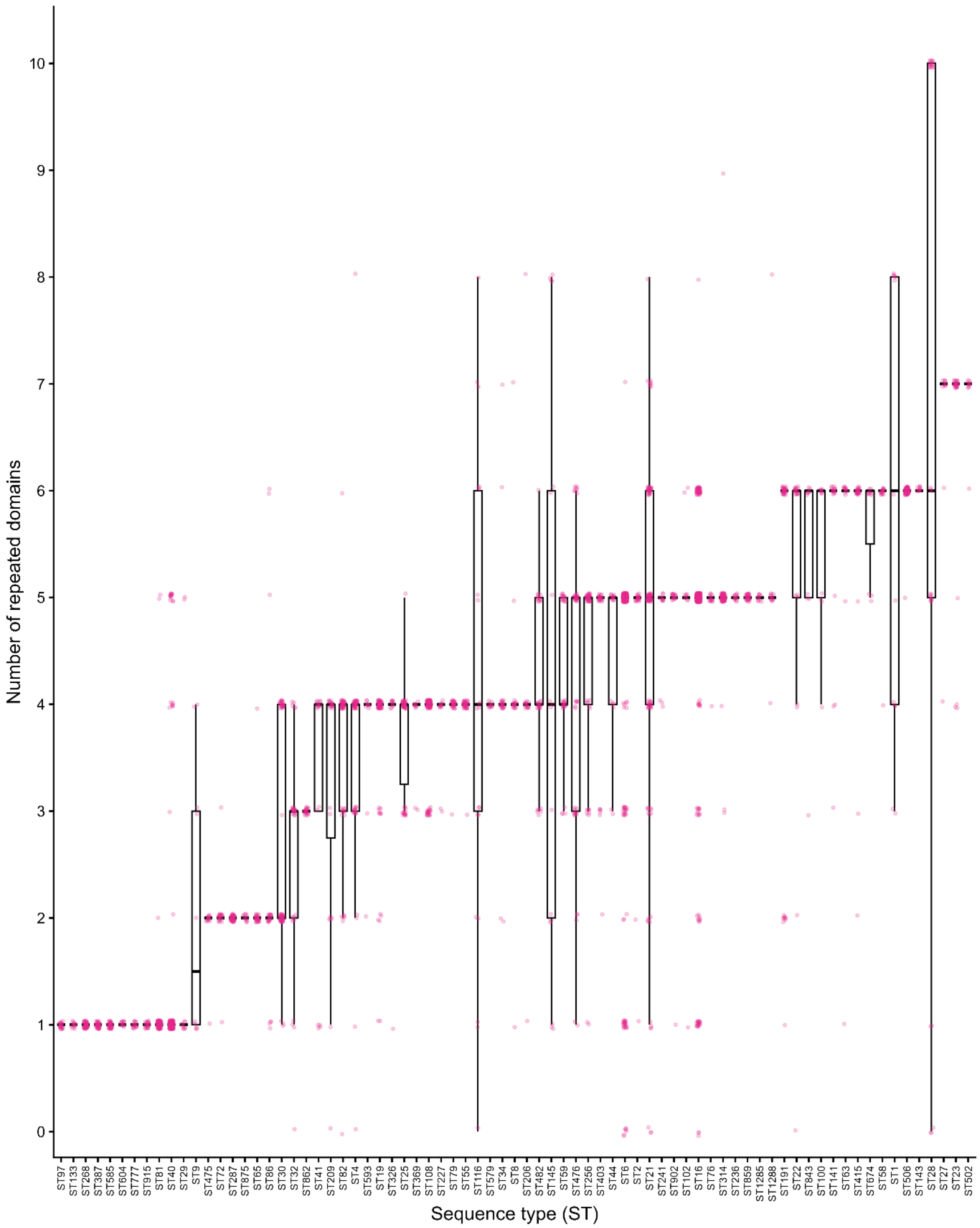


**Supplementary Fig. S5. Number of FraA CLIPPER domains across *E. faecalis* sequence types (ST).** Variation in CLIPPER domain number across sequence types. The number of CLIPPER domains per protein is shown for the 75 STs represented by at least 10 genomes encoding a full-length protein, ordered by increasing median domain number. Boxes indicate the interquartile range, horizontal lines indicate medians. Whiskers extend to the lowest and highest observations that fall within 1.5 times the interquartile range below the first quartile and above the third quartile, respectively. Pink points represent individual observations, jittered to reduce overlap.

Domain number differed significantly among STs (Kruskal–Wallis $\chi^2 = 2518.2$, df = 74, $p < 2.2 \times 10^{-16}$; $\varepsilon^2 = 0.717$).

Table S4. Bacterial strains used in this study

| Strain | Description | Source or reference |
|---|---|---|
| OG1RF | *Enterococcus faecalis*; $Rif^R$, $Fus^R$ | Dunny et al. (1978) |
| Δ*ebpA* | OG1RF *ebpA* deletion mutant | Nielsen et al. (2012) |
| Δ*ebpABC* | OG1RF *ebpABC* deletion mutant | Nielsen et al. (2012) |
| Δ*fraA* | OG1RF *fraA* (OG1RF_11764) deletion mutant | This work |
| V583 | Vancomycin-resistant *E. faecalis* bloodstream isolate | Sahm et al. (1989) |
| TTSHW-EF01 | *E. faecalis* wound infection isolate | This work |
| TTSHW-EF02 | *E. faecalis* wound infection isolate | This work |
| TTSHW-EF03 | *E. faecalis* wound infection isolate | This work |
| TTSHW-EF05 | *E. faecalis* wound infection isolate | Ch'ng et al. (2022) |
| TTSHW-EF07 | *E. faecalis* wound infection isolate | This work |
| TTSHW-EF24 | *E. faecalis* wound infection isolate | Ch'ng et al. (2022) |
| TTSHW-EF34 | *E. faecalis* wound infection isolate | This work |

Clinical isolates were collected from patients with wound infections at Tan Tock Seng Hospital, Singapore, under the supervision of T. M. S. Barkham (Department of Laboratory Medicine) and identified as *E. faecalis* by mass spectrometry. $Rif^R$, rifampicin resistant; $Fus^R$, fusidic acid resistant; Δ, deletion.

Table S5. Primers used in this study

| Primer | Application | Sequence (5'–3') |
|---|---|---|
| M13 F | pGCP213 screening | GTAAAACGACGGCCAGT |
| M13 R | pGCP213 screening | CAGGAAACAGCTATGAC |
| 11764_del1 | *fraA* deletion construction | AAGGAATTCGCAATAGCAAATGTAGCTAAGC |
| 11764_del2 | *fraA* deletion construction | CAATCGATTTTGTTTACATTCTCCTTCGTTCTTTTC |
| 11764_del3 | *fraA* deletion construction | GAACGAAGGAGAATGTAAACAAAATCGATTGTTTTAGAAC |
| 11764_del4 | *fraA* deletion construction | AAGAAGCTTAAAAACAATTGGCTAAACC |
| 11764 screen F | *fraA* deletion screening | ATTCACGATTTTTTGTTGAAATATCG |
| 11764 screen R | *fraA* deletion screening | TATCCGTAAAAAACACTAGAGC |
| 11764_FW | CLIPPER repeat-number screening | CAGCCCGGGGGATCCATGAATAAAGCAGTTAAAAATTTTG |
| 11764_RV | CLIPPER repeat-number screening | CGCTCTAGAACTAGTTTACTCCTCTTTTTCTTTGTTTCTC |

The gene *fraA* corresponds to OG1RF_11764. Primer names retain their original laboratory identifiers.
